# Widespread genomic heterogeneity at the type II NAD(P)H dehydrogenase locus predisposes *Cryptosporidium* to clofazimine resistance

**DOI:** 10.1101/2025.10.07.680968

**Authors:** Gracyn Y. Buenconsejo, Sebastian Shaw, Rui Xiao, Aurelia Balestra, Keenan M. O’Dea, Peng Jiang, Bingjie Xu, Dongqiang Wang, Guan Zhu, Daniel P. Beiting, Boris Striepen

**Author notes:** These authors contributed equally (Gracyn Y. Buenconsejo, Sebastian Shaw).

## Abstract

The parasite *Cryptosporidium* is a leading cause of life-threatening diarrheal disease, and effective treatment is not available. The discovery of potent anti-*Cryptosporidium* activity of clofazimine, offered the opportunity to repurpose a drug already used to treat leprosy and tuberculosis. However, clofazimine failed in a human trial, which was attributed to poor bioavailability. Here, we observed differential susceptibility among parasite isolates which we exploit to map the mode of action to type II NADH dehydrogenase (NDH2) in an unbiased genetic cross. Targeted genetic ablation of NDH2 resulted in profound clofazimine resistance, and biochemical studies demonstrated NDH2 mediated electron transfer to clofazimine. Through genomic analyses we uncovered heterogeneity at the NDH2 locus for *C. parvum* and *C. hominis* and widespread carriage of a conserved attenuated allele across multiple continents. This heterogeneity allows parasites genomically linked through frequent sexual recombination to adjust to changing NDH2 requirements and predisposes *Cryptosporidium* to evade clofazimine treatment.

## Introduction

The apicomplexan parasite *Cryptosporidium* is an important cause of intestinal disease in a variety of epidemiological settings. The transmissive oocyst stage is inherently resistant to water chlorination and waterborne outbreaks occur regularly in the United States despite advanced water treatment in place. *Cryptosporidium* has long been recognized as an life-threatening opportunistic infection in HIV/AIDS patients, causing watery diarrhea and wasting associated with poor prognosis^1^. Various conditions resulting in reduced cellular immune function including certain cancers, solid organ transplantation and the accompanying immunosuppressive therapy, and multiple primary genetic defects similarly predispose to life-threatening cryptosporidiosis ^2^. Despite decades of effort, effective treatment is still unavailable and the clinical management of cryptosporidiosis remains very difficult in these patients ^3^. More recently *Cryptosporidium* was also identified as a leading global cause of severe diarrhea and associated deaths in immunocompetent young children, particularly those experiencing malnutrition^4^. In a vicious circle, cryptosporidiosis itself predisposes children to malnutrition and stunting^5^. In the absence of a vaccine, effective drugs are urgently needed for the treatment of this large pediatric population as well ^6–8^. Motivated by this need academic and industry laboratories conducted multiple large-scale screens to identify new anti-parasitic compounds^6,9^. Acknowledging the economic challenge of drug development for a largely resource poor target population, several of these efforts attempted to leverage prior investments through repurposing of established drugs, screening leads, or cherry picked compound libraries ^10–12^ often focused on antimalarials.

Among the findings of these studies was the discovery of potent anti-*Cryptosporidium* activity of clofazimine ^11^, which was widely welcomed, as this drug is already in clinical use for decades and is relatively inexpensive to produce. Clofazimine is a riminophenazine derivative (Fig. 1a) developed in the 1950s for the treatment of tuberculosis that was initially sidelined in favor of more potent antibiotics^13^. However, the drug has been in widespread use for the treatment of leprosy, where it is used routinely in combination therapy with rifampin and dapsone^14^. Recently multi-drug resistance has spurred a renewed interest in clofazimine for treating tuberculosis. Following clinical trials ^15^ the WHO now suggests the use of a 6-month treatment regimen composed of bedaquiline, delamanid, linezolid, levofloxacin, and clofazimine in multidrug-resistant/rifampicin-resistant tuberculosis patients with or without fluoroquinolone resistance ^16^. Important questions remain in the tuberculosis field about clofazimine’s safety and side effects, its potential for antimicrobial resistance, and related to both of these questions is the lack of conclusive understanding of its mode of action ^17^. The molecule is highly lipophilic and therefore thought to exert its activity at the level of cellular membranes. Among the mechanisms that are debated are: binding to bacterial DNA and inhibiting replication^18^, interference with bacterial redox and energy balance due to competition with menaquinone as an electron acceptor for type 2 NAD(P)H dehydrogenase (NDH2) which is part of the bacterial respiratory chain ^19,20^, or interference with bacterial potassium transport either directly or mediated through phospholipid metabolism ^21–23^. Clofazimine resistance has been reported clinically and in experimental selection in *Mycobacterium tuberculosis*, but without clear genetic clues as to the target and mode of action of the drug^17^.

**Figure 1:**
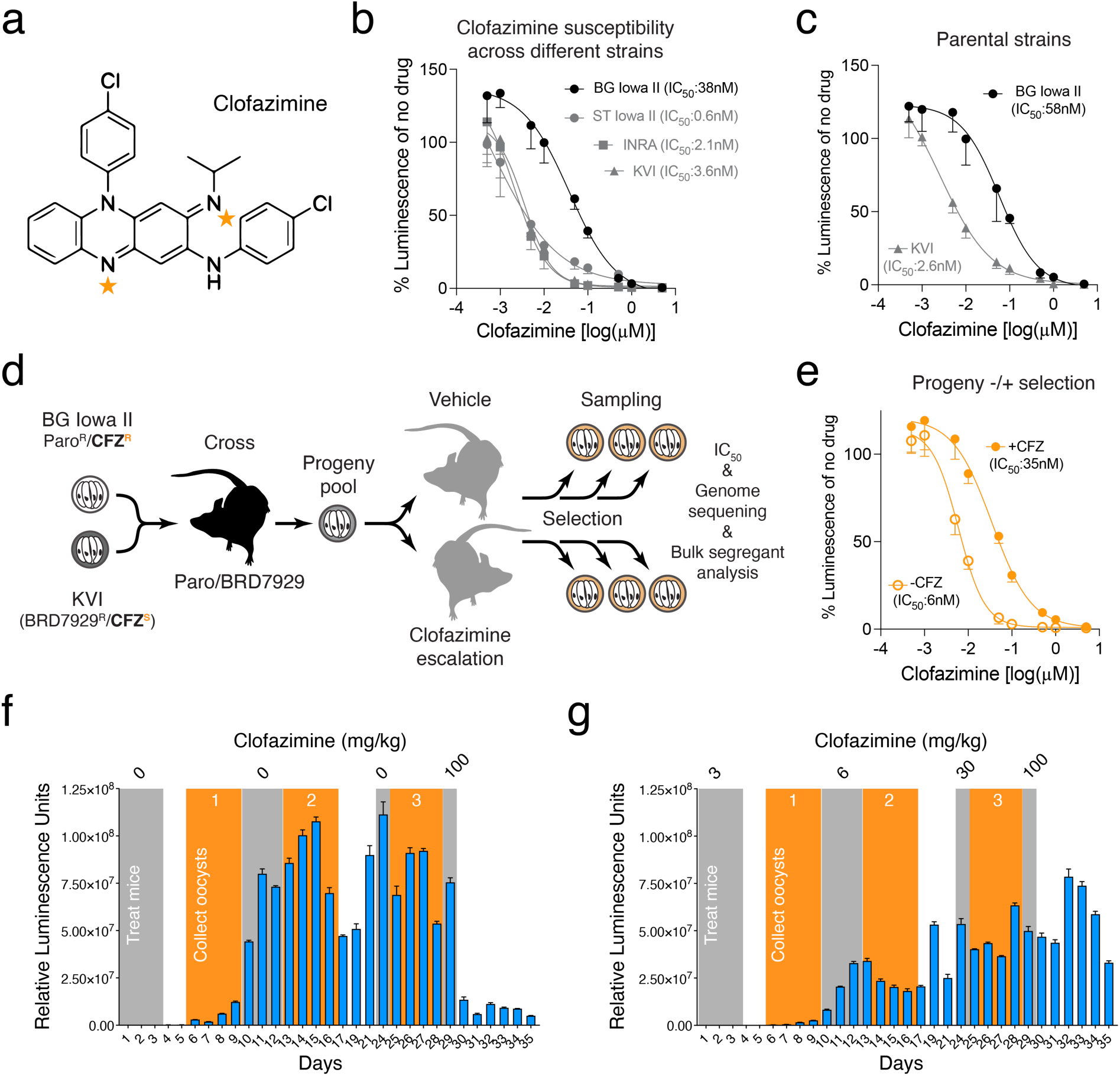
A genetic cross of *C. parvum* strains differing in clofazimine (CFZ) susceptibility. (**a**) Chemical structure of clofazimine. Tangerine stars indicate atoms positioned to accept electrons during enzymatic reduction. (**b** and **c**) Parasite growth in HCT-8 tissue culture assessed by measuring luminescence in the presence of the indicated concentrations of clofazimine. Data represent mean and SD of n=5 biological replicates. (**d**) Experimental set up of genetic cross between BG Iowa II (CFZ-resistant) and KVI (CFZ-susceptible). Recombinant progeny was used to infect mice treated with vehicle or escalating doses of CFZ. Oocysts were sampled throughout the selection for IC_50_determination and genome sequencing. (**e**) HCT-8 culture clofazimine susceptibility assay of cross progeny treated with vehicle or drug (mean and SD of n=5 biological replicates). (**f** and **g**) Parasite burden was measured by following fecal luciferase activity for progeny treated with vehicle (**f**) or clofazimine (**g**); *n* = 3 mice per cross. Gray boxes indicate treatment windows. Orange boxes indicate times oocyst were collected.

Most disappointingly, clofazimine failed in human trial for the treatment of cryptosporidiosis. The trial was designed as a randomized, double-blinded, placebo-controlled study in HIV infected adults suffering from cryptosporidiosis ^24^. This was a difficult study, challenged by the very poor health of the participants, difficulties in recruitment, and a resulting lack of randomization. Beyond the lack of efficacy, the study also observed lower than anticipated serum levels for clofazimine in the *Cryptosporidium* infected treatment group, which was attributed to the severity of diarrheal diseases in those participants. Further studies let the authors to conclude that lower than required bioavailability of the drug ^25,26^ was likely responsible for the observed lack of efficacy.

Using a combination of forward and reverse genetic experimentation, we map clofazimine sensitivity of *Cryptosporidium* to its NDH2 gene and population genomic studies reveal widespread genomic heterogenicity at this specific locus in both *C. parvum* and *C. hominis* genomes across the globe. These genetic findings have important consequences for the rapid emergence of resistance and provide additional clues to the interpretation of the clinical failure of clofazimine for the treatment of cryptosporidiosis. They also suggest an unconventional role of NDH2 outside of the mitochondrial respiratory chain, one that appears to benefit from the ability to maintain and modulate a pseudo-diploid stage in a haploid organism through frequent sexual recombination.

## Results

### Clofazimine susceptibility varies across different *C. parvum* strains

Clofazimine failed in human trial despite preclinical promise, and this has been linked to poor bioavailability of the drug in patients with severe diarrhea ^25,26^. However, the initial report describing the anti-*Cryptosporidium* activity of clofazimine noted lower potency against *C. hominis* in culture assays. These differences in clofazimine susceptibility as well as preliminary observations on two *C. parvum* strains, led us to consider the possibility of strain-specific, heritable differences in clofazimine susceptibility. To test this more rigorously we established the half-maximal inhibitory concentration (IC_50_) for clofazimine for multiple different *C. parvum* strains. We used ST Iowa II, originally obtained from the Sterling laboratory at the University of Arizona and used in the original drug screen, BG Iowa II (a closely related strain propagated by Bunchgrass Farms and widely used for laboratory experiments), and INRA initially isolated in France. All three are IIa genotypes derived from cattle, in contrast, KVI is a IId strain recently isolated from an infected lamb in Israel ^27^. All four strains were engineered to express nanoluciferase, and we measured parasite growth over 48h in a human ileocecal adenocarcinoma (HCT-8) cell culture ^28^ and performed nine-point dose-response assays ranging from 0.5nM to 5µM with half-log _10_ steps (Fig. 1b, all values normalized to vehicle control for each strain). KVI, INRA and Sterling showed similar susceptibility (IC_50_=3.6nM, 2.1nM and 0.6nM, with 95% confidence intervals (CI) of 2.6 to 4.3, 0.1 to 4.1, and ∞ to 3, respectively) comparable to that initially reported by Love et al., while BG Iowa II was about 10-20 times more resistant (IC_50_=38nM; 95% CI=27 to 52) akin to the earlier measurement for *C. hominis*^11^.

### Selecting for clofazimine resistance in a genetic cross

We recently developed genetic crosses to study *Cryptosporidium* ^27,29^ and wondered whether we might be able to exploit this differential susceptibility to map the mechanism of action for clofazimine in this parasite in an unbiased forward genetic fashion. We chose Bunchgrass and KVI as crossing parents as they offer the largest number of distinguishing single nucleotide polymorphisms (SNPs) and validated the susceptibility differential in strains engineered with marker cassettes suitable for a cross (Neo for BG^28^ and mutated phenylalanyl tRNA synthase ^29,30^ for KVI, Fig. 1c). Fig. 1d outlines the design of the cross, interferon-γ knockout (*ifnγ^−/−^*) mice were infected with both parental parasite strains and treated with BRD7929 and paromomycin to select for recombinant progeny carrying both resistance markers. Oocysts of this progeny were collected and used to infect two new groups of mice, one was treated with escalating doses of clofazimine, the control with vehicle alone, and parasite burden was measured by fecal luciferase for 35 days (Fig. 1f and g). This treatment regime transiently repressed parasite growth but did not cure the mice as infection rebounded each time. We collected feces in the recovery periods (orange boxes) following each treatment (gray boxes) with drug or vehicle and isolated oocyst pools. We measured the impact of this selection scheme on clofazimine susceptibility in an in vitro dose-response assays for the final selected progeny pools (derived from days 25-28 post-infection following treatment with 30 mg/kg clofazimine or vehicle). The clofazimine-selected progeny showed marked reduction in susceptibility (IC_50_=35nM; 95% CI=26 to 47) when compared to the vehicle-treated progeny (IC_50_=6nM; 95% CI=5 to 7, Fig. 1e) suggesting selection for resistance.

### A single genomic locus is linked to clofazimine susceptibility

Genomic DNA was extracted from 5×10^6^ oocysts of each of the three pools collected for clofazimine selected and vehicle treated progeny populations, and we carried out high-throughput sequencing to generate robust genome coverage (134-228-fold). Reads were aligned to the *C. parvum* genome and SNPs were called for the 4700 positions discriminating the parental strains as detailed in the Methods section. Fig. 2a shows allele frequencies across all eight chromosomes compared to the telomere-to-telomere *C. parvum* BG Iowa II genome^31^, 1 indicates exclusive BG Iowa II and 0 exclusive KVI inheritance. Clofazimine selected populations are shown in red, vehicle in blue, and later timepoints are shown in darker shades. We noted multiple peaks indicating preferred inheritance from one of the parents. These included the loci of the two selectable markers on chromosome 3 and 5, and loci on chromosomes 2, 6, and 7 associated with enhanced virulence and persistence of the KVI parent (please refer to our recent publication ^27^ for detail on these loci). However, only chromosome 7 showed differences when comparing clofazimine treated and untreated samples (Fig. 2b, each individual SNP is represented by a dot). In the absence of treatment, KVI alleles dominated due to the virulence locus on this chromosome, upon treatment BG alleles were heavily enriched pointing to preferred inheritance from the more clofazimine resistant parent. We next conducted bulk segregant analysis ^27,32^ to detect and measure genetic linkage. A single narrowly defined quantitative trait locus on chromosome 7 emerged, the statistical support for this locus increased with each round of treatment and dose escalation, with the most highly significant SNP exceeding a final G-value of 300 (Fig. 2c and d).

**Figure 2:**
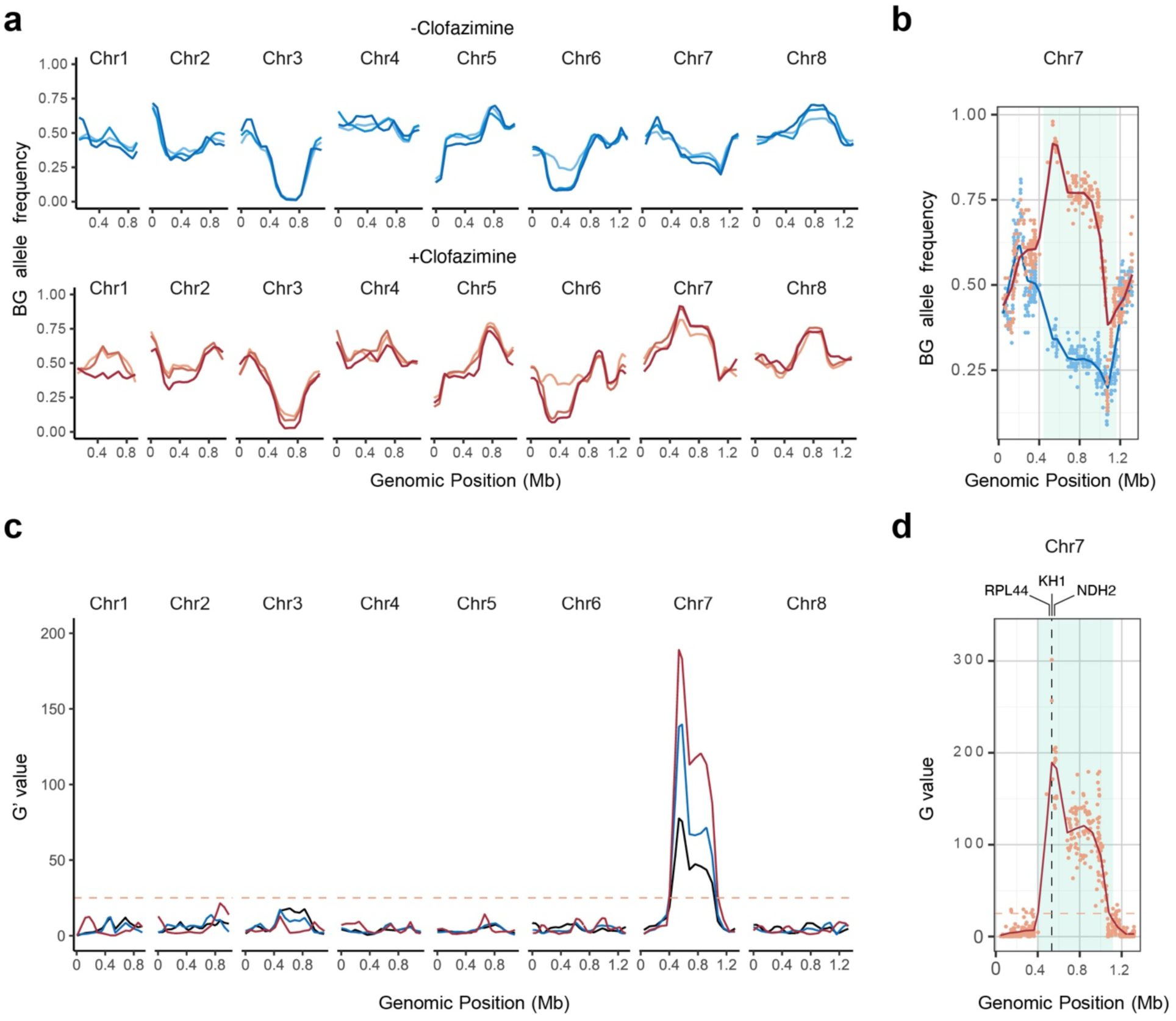
Clofazimine susceptibility is associated with a single locus on chromosome 7. (**a**) Allele frequencies of SNPs distinguishing the parental strains used in the cross for all eight *C. parvum* chromosomes. Each line represents a progeny pool, those shown in blue were treated with vehicle, those in red with clofazimine. Color shades from light to dark indicate the 3 individual pools that were collected, sequenced and analyzed. (**b**) Allele frequencies of chromosome 7 for the third and final pool collected. Blue = vehicle, red = clofazimine. (**c**) Whole genome G’-values for genetic linkage. Lines are the weighted moving averages for the G′-values, and the significance threshold is shown as an orange dashed line. Dark blue = pool 1, light blue = pool 2 and red = pool 3. (**d**) Highly significant QTL on chromosome 7. The G values of pool 3 for each individual SNP are shown as dots, the line is the weighted moving averages for the G′-values, and the significance threshold is shown as an orange dashed line. The 95th percentile of the QTL is shown in light blue.

### Resistance is linked to a two base pair deletion in the type 2 NAD(P)H dehydrogenase gene

The highest scoring SNP was found in gene cgd7_1890 resulting in a valine instead of an isoleucine in a KH1 domain-containing putative RNA-binding protein (Fig. 3a). This represents a conservative substitution, and while KH1 domain RNA binding proteins can play roles in drug resistance in some cancers^33^, they have not been previously associated with clofazimine resistance. We were thus intrigued to find NDH2 encoded by the next gene downstream of the SNP (cgd7_1900). NDH2 is one of the candidate mechanisms of clofazimine action in *Mycobacterium*^19^. However, the initial comparison of the published parental genomes showed identical NDH2 sequences in both strains. Our bulk segregant analysis used SNPs to detect QTLs. We considered that other variations may impact on drug susceptibility and therefore manually inspected the alignment of raw reads for this section. This revealed a previously unrecognized INDEL, the deletion of two adenines (ΔAA) at positions 81 and 82 of the open reading frame in 100% of all reads from clofazimine-selected parasites, while only 4.5% of all reads from the vehicle-treated parasites showed a deletion at this position. Alignment of the parental genomes detected the ΔAA allele in both parents, with higher frequency in the more resistant strain (Fig. 3b and 3c).

**Figure 3:**
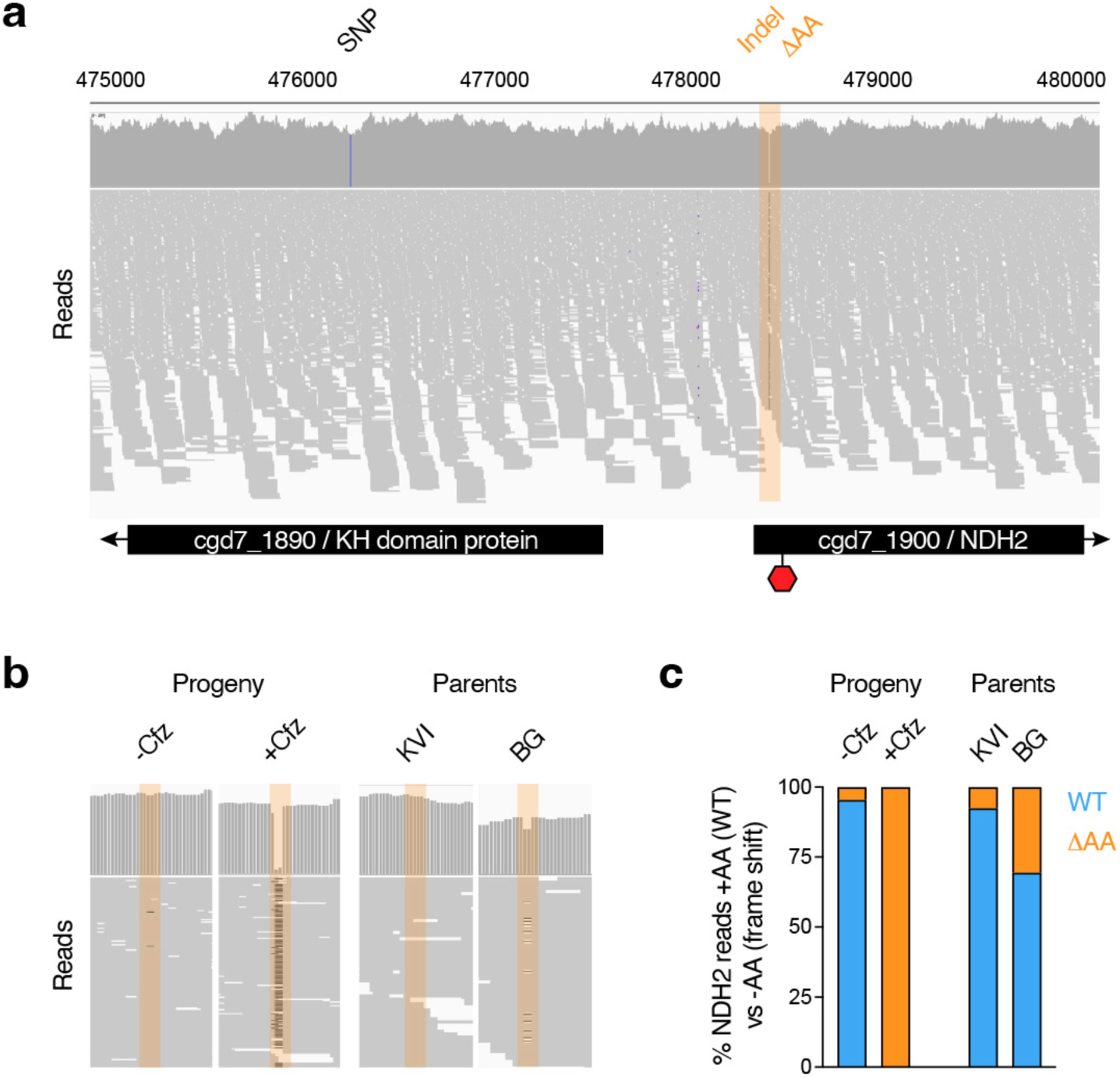
Clofazimine selected progeny carry an indel at the 5’ end of the NDH2 gene. (**a**) Illumina sequence read alignment of the clofazimine selected third progeny pool. Blue bar in gene cgd7_1890 indicates the SNP with the highest G value identified by bulk segregant analysis. Region highlighted in orange indicates the indel in the NDH2 gene that introduces a frameshift and a premature stop codon at the indicated position. (**b)** Indel frequency comparison across the progeny either selected or not selected with clofazimine and the parental strains. (**c**) Quantification of the indel frequency represented in (**b**).

### Genetic ablation of NHD2 confers high level clofazimine resistance

The ΔAA deletion is predicted to result in a frame shift of the NDH2 coding sequence and early termination of the protein after only 47 of its 568 predicted amino acids. We therefore hypothesized resistance to be the consequence of loss of NDH2 activity, rather than a SNP in KH1. To directly test this, we disrupted the NDH2 gene through Cas9 directed insertion of the Nluc/Neo marker in BG Iowa II parasites using paromomycin selection (Fig. 4a). Transgenic parasites were readily obtained, and PCR mapping verified appropriate insertion (Fig. 4b), and we conclude NDH2 to be dispensable. Next, we compared the sensitivity to clofazimine of the deletion mutant to wildtype BG or KVI parasites in tissue culture growth assays and found the mutant to be highly drug resistant with an IC_50_ of 4.1µM (Fig. 4c). Based on these findings we reasoned that disruption of the NDH2 locus should be selectable by clofazimine treatment. KVI strain sporozoites were electroporated with a marker-less targeting vector encoding Nluc and tdNeonGreen fluorescent protein along with a plasmid for the expression of Cas9 and a guide RNA targeting NDH2 (Fig. 4d). Mice infected with these sporozoites were treated for 7 days with 100 mg/kg/day clofazimine starting 24h post-infection. Activity of the Nluc transgene was detected in the feces of these mice on day four following transfection and continued to rapidly increase (Fig. 4f). PCR analysis of genomic DNA demonstrated transgene insertion and complete loss of the wildtype locus (Fig. 4e). Oocysts subjected to flow cytometry showed uniformly bright fluorescence and when used to infect HCT-8 cultures all parasite stages were Neon Green positive when observed by fluorescence microscopy (Fig. 4g and 4h). We conclude that loss of NDH2 confers resistance to clofazimine, and that clofazimine offers a new selection principle for *Cryptosporidium* transgenesis.

**Figure 4:**
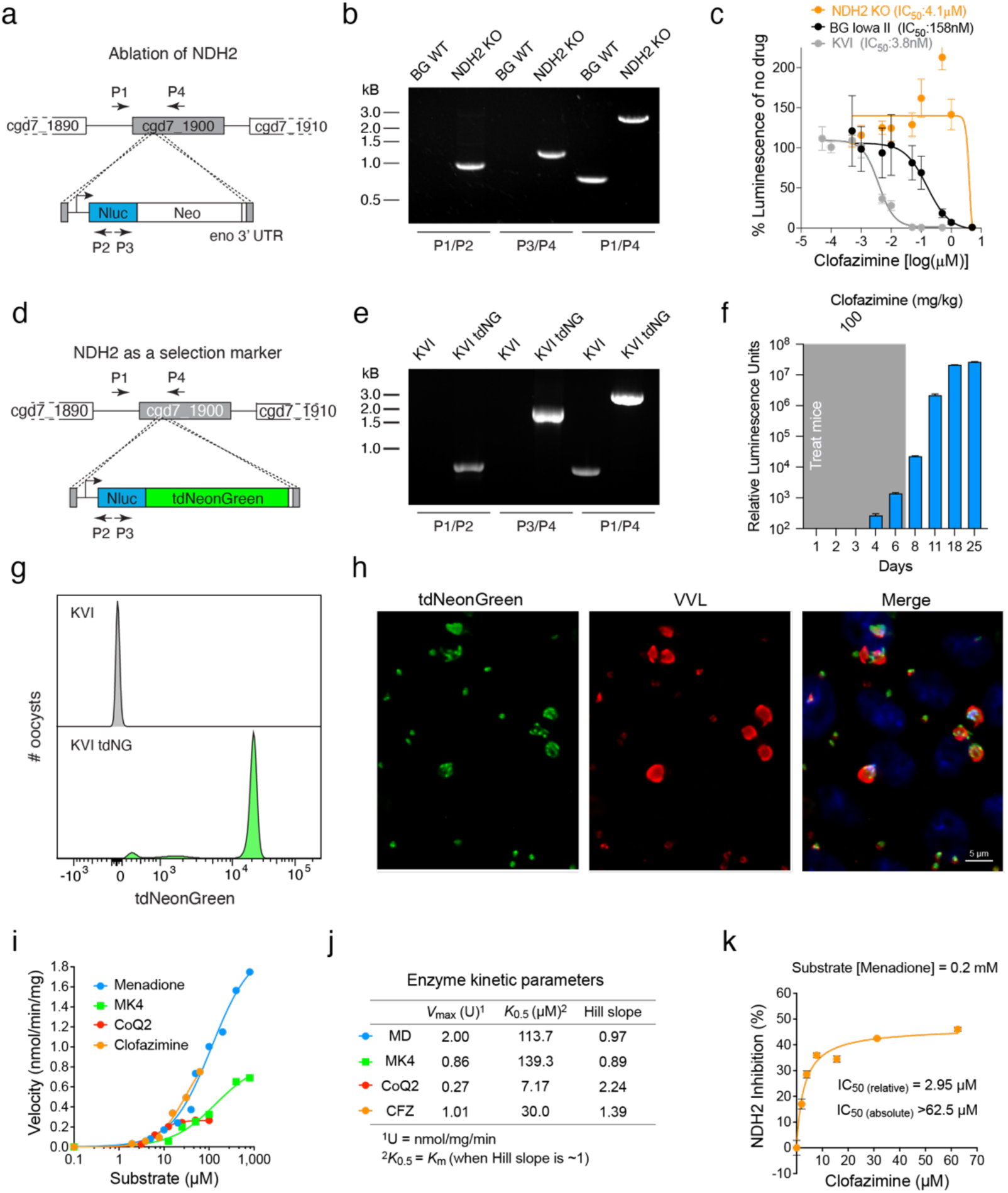
NDH2 ablation results in high level clofazimine resistance. (**a**) Genetic ablation of NDH2 by Cas9 mediated marker insertion. P1 – P4 indicate positions of primers used for PCR mapping (**b**) of the mutant and parental wildtype (WT) strains. (**c**) Clofazimine IC_50_ determination in HCT-8 culture of the NDH2 knockout and wildtype strains used in the cross. Data are mean and SD of n=5 biological replicates. (**d**) Strategy and constructs used to test whether NDH2 could be used as a selection marker for *C. parvum*. P1 – P4 indicate positions of primers used for PCR mapping (**e**) of the resulting transgenics compared to KVI wildtype. (**f**) Luciferase activity of parasites transfected as shown in (**d**) and selected with clofazimine. (**g**) Oocyst of clofazimine selected transgenic were purified from the feces and subjected to flow cytometry (gray: KVI WT; green: KVI tdNeonGreen) or used to infect HCT-8 cells (**h**) and observed by fluorescence microscopy at 48h of culture. Note bright green fluorescence in flow cytometry histogram and micrograph (gray: Hoechst; green: tdNeonGreen; red: VVL). Scale bar, 5 μm. (**i**) Spectrophotometric assay of CpNDH2-WT showing NADH-dependent reduction of menadione (MD), menaquinone-4 (MK4), ubiquinone-2 (CoQ2), and clofazimine (CFZ). Michaelis–Menten kinetics were observed with MD and MK4, while CoQ2 and CFZ showed positive cooperativity. Catalytic efficiency ranked MD > MK4 > CoQ2; CFZ activity was comparable to quinones. (**j**) Kinetic parameters (*V*_max_, *K*_m_ or *K*_0.5_, Hill slope) derived from nonlinear regression are summarized in the table. (**k**) Clofazimine inhibits CpNDH2-mediated MD reduction in a dose-dependent manner (relative *IC*_50_ ≈ 2.95 μM; absolute *IC*_50_ > 62.5 μM). The results support competitive reduction of CFZ by CpNDH2. Primers used for genotyping in (**b**) and (**e**) can be found in Supplementary Data 3.

### Recombinant *C. parvum* NDH2 recognizes clofazimine as substrate

Biochemical studies in bacteria have suggested that clofazimine may compete with natural quinones for electron transfer by NDH2 (the two critical nitrogen positions are highlighted in Fig. 1a), with subsequent spontaneous reactions giving rise to reactive oxygen species that ultimately damage the cell^19^. To test this for *Cryptosporidium*, we expressed *C. parvum* NDH2 in *Escherichia coli* and purified the recombinant maltose-binding protein fusion in intact and cleaved form (Supplementary Fig.1a). A spectrophotometric assay was established to measure the ability of recombinant enzyme to transfer electrons from NADH to various quinone substrates ^34–36^. Initial assessment showed that both intact and cleaved forms could catalyze the electron transfer from NADH to menadione with the same efficiency (Supplementary Fig.1b and 1c). We observed activity with menadione (MD), menaquinone-4 (MK4), and ubiquinone-2 (CoQ2) as substrates, with low-micromolar *K*_m_ values (or *K*_0.5_ for CoQ2, which shows positive cooperativity, Fig. 4i and j). The enzyme showed highest activity with menadione (*K*_m_ = 113.7 μM; *V*_max_ = 2.0 U, U = nmol/min/mg), and greater activity with short-chain menaquinone (*K*_m_ = 139.3 μM; *V*_max_ = 0.96 U on MK4) than with short-chain ubiquinone (*K*_m_ = 7.17 μM; *V*_max_ = 0.27 U on CoQ2). These *K*_m_ (or *K*_0.5_) values are comparable to those reported for NDH2 from other organisms, including *Saccharomyces cerevisiae* (15.2 and 7.9 μM on CoQ1 and CoQ2, respectively) ^37^ and *Caldalkalibacillus thermarum* (34.0 μM on MD)^38^. *C. parvum* NDH2 also reduced clofazimine, showing activity comparable to that observed for quinones (*K*_m_ = 30.0 μM; *V*_max_ = 1.0 U). Moreover, clofazimine competed with native substrates, with a relative IC_50_ of 2.95 μM for menadione reduction (Fig. 4k). Collectively, these biochemical experiments confirmed that *C. parvum* NDH2 indeed has type 2 NDH activity, and that clofazimine can be reduced by this enzyme.

### NDH2 localizes to the inner membrane complex and not the mitosome

NDH2 typically acts in the respiratory chain and in bacteria is associated with the cell membrane. In eukaryotes including related apicomplexan parasites it is a mitochondrial protein. The *C. parvum* mitochondrion has lost its genome and much of its respiratory metabolism ^39^, this small typically round organelle known as the mitosome is found in close proximity to the nucleus ^40^. We were thus surprised to consistently observe NDH2-HA staining as a line close to the surface of parasites, regardless of whether we expressed a transgene or tagged the native locus. High resolution expansion microscopy of infected HCT-8 cultures showed labeling in all intracellular stages of the parasite (Fig. 5a). NDH2 labeling outlined extracellular merozoites and male gametes beginning during the intracellular assembly, and in non-dividing parasites appeared as a sharply delineated cap underlying the membrane facing away from the host cell (Fig. 5a, single arrowhead indicates this cap in a female gamete). Higher magnification reveals this labeling to coincide with the inward facing membrane of the inner membrane complex (IMC, Fig. 5b)^41^. We confirmed IMC assignment by co-labeling with an antibody to the conserved alveolin domain of IMC proteins^42,43^ (Fig. 5c this epitope did not tolerate the expansion protocol). For comparison we also introduced an epitope tag into cgd8_380 which encodes malate oxidoreductase, a presumptive mitosomal protein ^44^. For this protein we indeed observed localization to a small organelle close to the nucleus (Fig. 5d). We find labeling in all stages with exception of male gametes matching recent findings by Li and colleagues for alternative oxidase ^45^. We conclude that NDH2 in *C. parvum* is not a mitochondrial protein but is localized to the membrane of the IMC facing the parasite cytoplasm.

**Figure 5:**
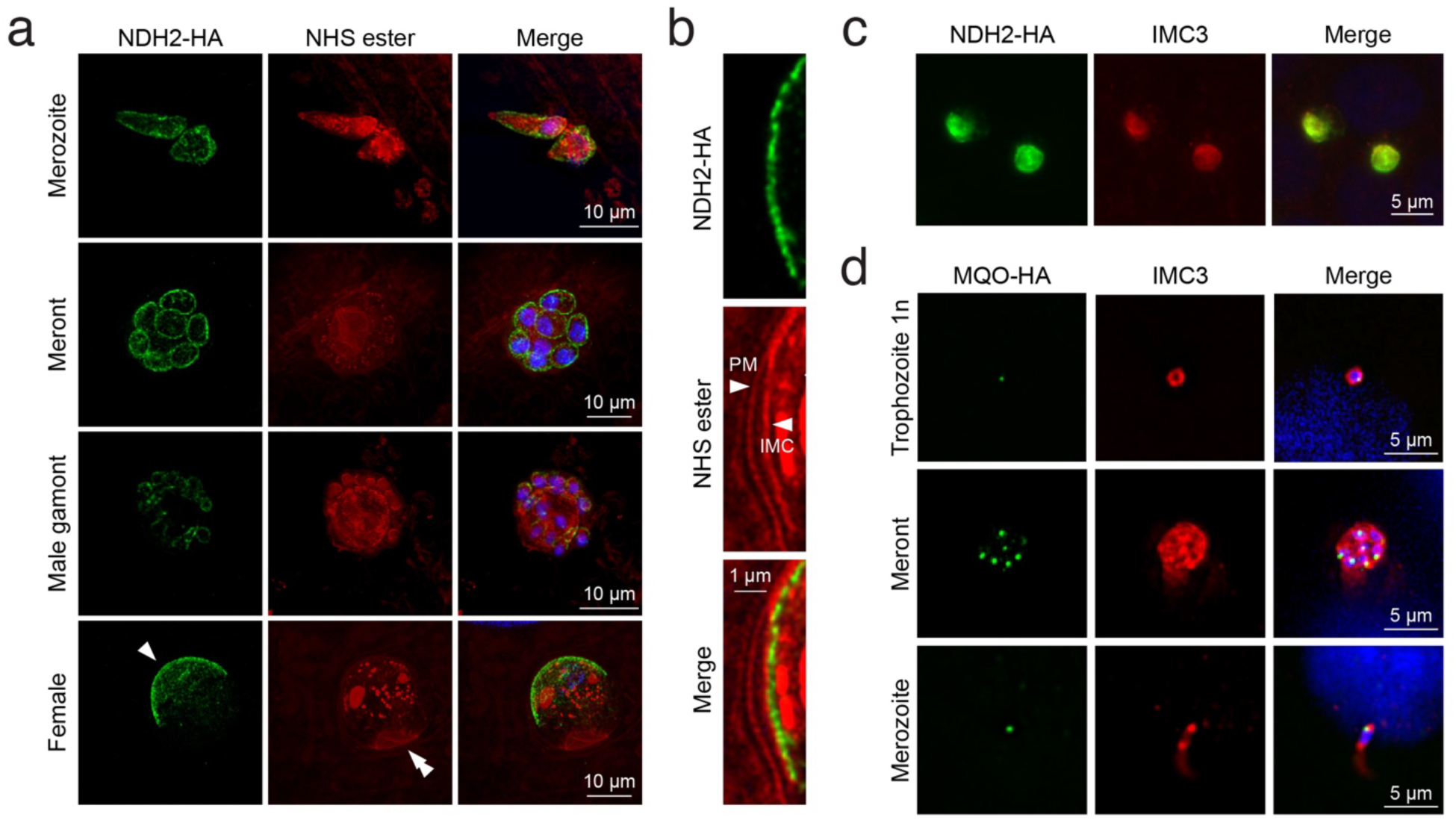
NDH2 is localized to the inner membrane complex. (**a**) Expansion microscopy of HCT-8 cultures infected with NDH2-HA parasites showing protein localization throughout the life cycle. HA in green; NHS-ester in red; Hoechst in blue; scale bar = 10 μm. Single arrowhead NDH2 at the IMC, double arrowhead highlights host-parasite interface at the base of the intracellular parasites in side view. (**b**) Higher magnification detail of the female parasite shown in (**a**) PM = parasite membrane; IMC = inner membrane complex; HA in green; NHS-ester in red; Hoechst in blue; scale bar = 1 μm. Note labeling on the cytoplasmic side of the inner face of the IMC (**c**) Widefield image of NDH2-HA cells labeled antibodies to HA and IMC3 showing colocalization. HA = green; IMC3 = red; scale bar = 5μm. (**d**) Widefield image of MQA-HA parasites labled with antibodies to HA and IMC3 showing punctate MQO localization consistent with targeting to the mitosome. HA = green; IMC3 = red; Hoechst = blue; scale bar = 5μm.

### Genomic heterogeneity at the NDH2 locus and the ΔAA allele are widespread

We were initially surprised to find that both parental strains used in the cross carried the ΔAA allele, albeit at different frequencies (30.5% in BG Iowa II and 7.4% in KVI, Fig. 3b and 3c). This motivated a broader analysis in which we analyzed publicly available whole genomic sequencing (WGS) data from 68 *C. parvum* and *C. hominis* sequencing read archives. These were selected for robust genome coverage (>30X mean depth, with mean mapping quality>=60) and broad geographic representation (Fig. 6a and 6b). Reads were aligned to the *C. parvum* reference genome to call variants (INDELs and SNPs with a rigorous quality score >=20 and depth>=20), and we scored the frequency of the ΔAA allele. Remarkably, the same ΔAA variant is detectable in the NDH2 genes of most genomes analyzed (91%) (Supplementary Data 1). In most strains at a frequency around 5-10% while BG Iowa II stands out (note that this genome was sequenced multiple times). We wondered whether this frequency may simply reflect a broader tendency of *Cryptosporidium* genomes to hypermutation and heterogeneity. As a control, we scored all high impact variations for NDH2 along with three essential and three dispensable genes across 38 *C. parvum* WGS SRAs of diverse geographic origin. Fig. 6c shows incidence of variation as a heatmap normalized into 10 bins for each gene’s coding sequence region to account for difference in length. The 5’ end of NDH2 clearly stands out, and inspection showed the ΔAA allele to account for all this variation.

**Figure 6:**
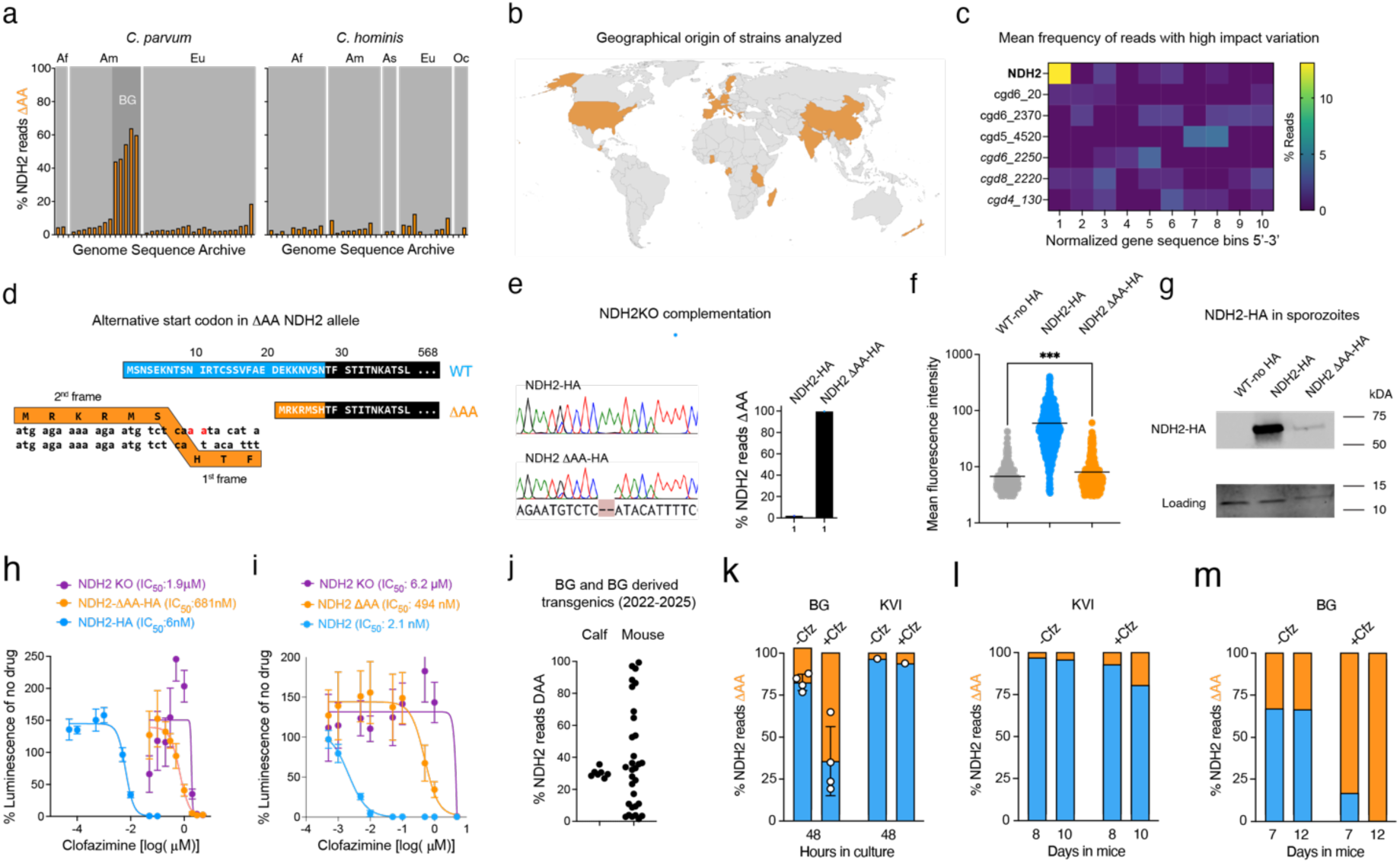
NDH2 heterogeneity is widespread and responds dynamically to drug pressure. (**a**) Analysis of ΔAA allele frequency across multiple publicly available whole genome sequences of *C. parvum* and *C. hominis* strains. Af = Africa; Am = America; Eu = Europe; As = Asia; Oc = Oceania; BG = Bunchgrass Iowa II. Filters were rigorously set to a minimum of 20 unique reads mapped at this location with minimum mapping quality of 60. See Methods section for detail and all data for specific genomes is shown in Supplementary Data 1 along with accession numbers. (**b**) Map showing the global origins of the samples analyzed in (**a**) in orange. (**c**) Heatmap representing high impact variations within the open reading frame of NDH2 and 3 dispensable genes (regular font) and 3 essential genes (italic font). (**d**) Reading frame and amino sequence at the start of NDH2 wildtype (blue) and potential alternative for the ΔAA allele (orange). (**e**) Sanger sequencing and amplicon sequencing of NADH2-HA and NDH2 ΔAA-HA transgenics derived by complementation of the NDH2KO. (**f**) Mean fluorescence of WT BG Iowa II parasites (n = 439) lacking the HA epitope and NHD2-KO parasites expressing NDH2-HA (n = 638) and NDH2 ΔAA-HA (n = 578) in HCT8 cultures labeled with antibody to HA. Statistical analysis was performed using one-way ANOVA and unpaired t test; P value = 0.0004. (**g**) Western blot of WT sporozoites lacking the HA epitope and NHD2-KO sporozoites expressing NDH2-HA or NDH2 ΔAA-HA. Histone 3 served as loading control. Apparent molecular weights of NDH2-HA and NDH2 ΔAA-HA are 72 and 70 kDa, respectively. (**h** and **i**) IC_50_ determination comparing the complemented strains and NDH2 KO (**h**) and the in situ HA-tagged strains homogenizing the NDH2 locus to WT or ΔAA allele (**i**). Data are mean and SD of n=5 biological replicates. (**j**) Amplicon sequencing measuring ΔAA allele frequency across five different batches of wildtype bunchgrass farm oocysts and 33 transgenic oocysts sampled from 2022 to 2025. (**k**-**m**) Effect of clofazimine treatment on ΔAA allele frequency of BG and KVI strain parasites in tissue culture (**k**) and in infected mice (**l** and **m**) measured by amplicon sequencing.

### The ΔAA allele attenuates but does not fully ablate NDH2 activity

Analysis of the NDH2 gene shows the ΔAA allele to break the reading frame, however, the use of an alternative start codon might result in a largely intact enzyme with an altered N-terminus (Fig. 6d). To better understand the impact of the ΔAA allele we engineered two pairs of parasite strains to differ at this specific position of the gene. These were constructed either by editing the native NDH2 locus to homogenously WT or ΔAA in the BG Iowa II strain, which incurred a small change in the protein sequence as well, or alternatively by complementing the knockout mutant with wildtype NDH2 or NDH2-ΔAA transgenes inserted into a neutral locus (see Supplementary Fig. 2 and Methods for detail). All genotypes were validated by PCR analysis and sequencing (Fig. 6e). Engineering these strains, we also introduced HA-epitope tags allowing to measure protein abundance. When imaging infected cultures by immunofluorescence, WT NDH2 is readily observed in all intracellular stages, the ΔAA signal is much lower, however, it is detectable above the level of an untagged control (Fig. 6f). Both proteins showed IMC localization. Western blot analysis of protein lysates generated from sporozoites showed a single band of the expected size for WT NDH2-HA and a much weaker, yet again detectable band for NDH2-ΔAA-HA (Fig. 6g). The predicted and measured apparent molecular weights of both proteins are near identical. Next, we conducted IC_50_ determinations in culture for both strain pairs in direct comparison to the NDH2-KO and found that sole expression of, or complementation with the WT allele produced clofazimine susceptibility with low nanomolar IC_50_s (2.1nM and 6nM, 95% CI=1.4 to 2.9 and 5.7 to 7.2). In contrast, the ΔAA allele conferred resistance in both backgrounds (in situ IC_50_=494nM; 95% CI=292 to 860; ectopic IC_50_=681nM; 95% CI=535 to 885), yet not to the full level of NDH2 deletion (Fig. 6h and 6i). We conclude that the ΔAA allele does not result in complete loss of NDH2 activity but severely attenuates its abundance leading to robust resistance.

### NDH2 allele frequency is dynamic and responds to environmental change

We wondered how dynamic the NDH2 locus might be and used amplicon sequencing to measure allele frequencies. We typically purchase BG Iowa II parasites from a commercial vendor and use them to engineer transgenic parasite. The vendor passages BG in immunocompetent calves, while we maintain parasites in *ifnγ^−/−^* mice. We measured the ΔAA allele frequency across BG samples from 2022 to 2025 and found a stable mean of 30.2 with a standard deviation of ±2.8, however when analyzing numerous transgenics derived from these parasites over this time frame we measure a very broad distribution (Fig. 6j). This suggests that the ΔAA frequency can change, and that the specific ratio might depend on the host environment.

To test this further we explored the impact of NDH2 heterogeneity on clofazimine susceptibility. HCT-8 cultures were infected with the BG Iowa II or KVI and grown in the presence or absence of 50 nM clofazimine. Treatment roughly doubled the ΔAA frequency in both strains over 48h (Fig. 6k). We also surveyed the allele frequence in infected mice. In the absence of drug treatment, the NDH2 allele frequency remained constant, in contrast, clofazimine treatment resulted in an increase of the ΔAA allele in both strains (reaching 100% for BG Iowa II strain on day 10, Fig. 6l and 6m). We conclude that NDH2 heterogeneity predisposes to selectable drug resistance, and that the ease and speed with which resistance is achieved may vary among strains, and hinge on the initial level of the ΔAA allele in a population.

## Discussion

Lack of effective treatment remains the crucial challenge to the clinical management of cryptosporidiosis, with consequences most grave for individuals with enhanced susceptibility due to immune deficiency or malnutrition ^3,6^. The last decade has seen tremendous progress due to the influx of funding and multiple technological advances. Leveraging the resources and expertise of the global malaria drug development effort was highly productive, and many of the strongest leads emerged from screening libraries enriched for compounds that had already shown promise against *Plasmodium falciparum ^12,30,46-48^*. While numerous antimalarials were developed over the last century, malaria parasites have also demonstrated a remarkable ability to evade them, multi-drug resistance is rampant and poses a very serious threat to control ^49^. A recent report of *Cryptosporidium* developing resistance to methionyl t-RNA synthetase inhibition ^50^ raises concern also for this parasite.

Our study suggests that clofazimine, one of the most promising candidates for the treatment of cryptosporidiosis failed due to a preexisting, yet previously undetected drug-resistant allele. We identified differential susceptibility among *C. parvum* strains, matching previous observations for the single *C. hominis* strain tested ^11^. Forward genetic mapping implicated the NDH2 locus, and biochemical experiments showed the ability of NDH2 to transfer electrons to clofazimine suggesting a prodrug activation mode of action also considered for mycobacteria ^19,20^. In *Mycobacterium* NDH2 KO did not yield drug resistance ^51^, however, interpretation of this result is complicated by the presence of three NDH type enzymes. In contrast, *Cryptosporidium* has a single NDH, and we found loss of this enzyme to result in high level resistance, and this loss is readily selectable by drug pressure. Importantly, a resistance conferring allele is already present and globally distributed in the absence of drug pressure. Low bioavailability of clofazimine due to severe diarrhea contributed to its clinical failure ^24–26^, and additional medicinal chemistry is likely to enhance the formulation and bioavailability of the drug ^52,53^. However, our animal experimentation shows that depending on the initial allele frequency, resistance can be attained over the course of a single infection, which casts serious doubt on the prospect of clofazimine as an anti-*Cryptosporidium* agent. Overall, this suggests that considering the resistance potential is an important step in future preclinical evaluation of anti-*Cryptosporidium* drugs.

NADH including type 2 NDH typically function as part of the respiratory chain transferring electrons to a lipophilic quinone acceptor in the bacterial plasma membrane, or in eukaryotes, the inner mitochondrial membrane. In mycobacteria the enzyme is dispensable, if fatty acids are not a main carbon source ^51^. In apicomplexan parasites NADH2 replaces the canonical complex I found in many other eukaryotes ^44,54^. *T. gondii* has two NDH2 enzymes that are both dispensable ^55^, the single *Plasmodium* NDH2 can be ablated with little consequence to blood stages, however development in the mosquito is blocked ^56,57^. In other apicomplexans NDH2 is a mitochondrial enzyme, while in *Cryptosporidium* it is localized to the IMC underlying the parasites plasma membrane. Interestingly, it appears confined to one face of the IMC. This was an initially surprising finding but matches the recent spatial proteomic assignment ^58^. In contrast, alternative oxidase and malate oxidoreductase, the two other redox enzymes thought to use a quinol electron carrier in *Cryptosporidium* are indeed mitochondrial proteins (^45^ and Fig. 5d). The *C. parvum* mitochondrion is highly reduced, it has lost its genome, the TCA cycle and most of the electron transport chain, and even those elements remaining appear dispensable ^45^.

The function of NDH2 at the IMC membrane is unknown, but its relocation out of the mitosome likely deemphasizes its importance to mitochondrial respiration. In the facultative intracellular pathogen *Listeria monocytogenes* NDH2 impacts redox balance and virulence independent of the respiratory chain to adjust to different host niches and metabolic environments ^59^. It is thus possible that NDH2 and ubiquinone might mitigate oxidative stress in a changing metabolic environment, and its main role may be to balance the cytoplasmic NAD/NADH pool. The changed localization of NDH2 could reflect a more outward facing role in attaining and modifying critical metabolites. Multiple recent studies have found *Cryptosporidium* to interact with host and microbiome derived metabolites in ways that profoundly impact parasite survival and growth ^60–63^. NDH2 could play a role in detoxifying detrimental metabolites ^64^. Lastly, redox pathways play key roles in antimicrobial restriction and broader immune signaling, and NDH2 activity may interact with and modulate these pathways ^65^.

Why is genomic heterogeneity at the NDH2 locus and the specific ΔAA allele conserved and widespread? We propose that heterogeneity may represent a metabolic rheostat and might carry the benefit of adaptability. When BG Iowa II parasites are passaged in calves, they carried a high frequence of the attenuated ΔAA allele and this remained constant over years. Upon introduction into mice this frequency varied significantly. What exactly drives this change is not known yet, but host metabolism, host nutrition (high fat milk replacer versus rodent chow), divergent microbiome composition, or differential immune pressure could all impact on parasite redox balance. Apicomplexa are widely seen as transcriptionally hard-wired with only limited metabolic flexibility ^66,67^. *Cryptosporidium* is unique among apicomplexans in that it undergoes obligate sex every two days resulting in a very high rates of rapid recombination ^68,69^. Combining this feature with genomic heterogeneity may allow the population in a single host to act as functional diploid with the ability to dial up or down a particular allele to dynamically adjust to change.

## Online Methods

### Parasites

*C. parvum* isolates and derived transgenics used in this study were obtained and genotyped as described ^70^. Original sources are: BG Iowa II, Bunch Grass Farms, Deary, Idaho; ST Iowa II, Dr. Reed, University of Arizona; INRA, Dr. Fabrice Laurent, INRAE and University of Tours, Nouzilly, France; KVI, Dr. Yasur-Landau, Beit Dagab, Israel.

### Generation of transgenic strains

Guide oligo nucleotides (Sigma-Aldrich) were introduced into the *C. parvum* Cas9/U6 plasmid ^28^ by restriction cloning (see reference ^71^ for guide design) and repair templates were constructed by Gibson assembly (New England Biolabs). Excysted sporozoites were transfected as described in^71^. Oligos used for genotyping can be found in Supplementary Data 3.

### Ablation of NDH2

The insert encodes Nluc followed by the neomycin phosphotransferase drug-selection marker and was knocked into the NDH2 locus to induce a knockout.

Guide: 7_1900_guide_F / 7_1900_guide_R

Repair template: NDH2KO_F / NDH2KO_R

### NDH2 as a selection marker

The insert encodes Nluc followed by a tdNeonGreen.

Guide: 7_1900_guide_F / 7_1900_guide_R

Repair template: mNG_Cfz_F / mNG_Cfz_R

### NDH2 in situ HA-epitope

The insert encodes a recodonized version of NDH2 (position 81 to stop codon) with a triple HA-epitope sequence followed by Nluc and the neomycin phosphotransferase drug-selection marker. 5’ homology arms between the NDH2-HA and the NDH2-ΔAA-HA strains differ. Note that the NDH2-ΔAA-HA strain has a slightly altered amino sequence at the beginning of the transgene (Fig. 6d).

NDH2-HA guide: 7_1900_guide_F / 7_1900_guide_R

NDH2-ΔAA-HA guide: NDH2_guide_2_F / NDH2_guide_2_R

NDH2-HA repair template: 7_1900_repair_+AA_F / 7_1900_repair_R

NDH2-ΔAA-HA repair template: NDH2_-AA_2_F / 7_1900_repair_R

### Ectopic expression of NDH2-HA

The inserts encode the last 113 bp of the pheRS gene (cgd3_3320, recodonized) including the mutation that confers resistance (L482V) to BRD7929. This short sequence is followed by the whole NDH2 gene cassette (including its own promoter, and a triple HA-epitope sequence and the enolase 3’UTR). The only difference between the repairs is the presence or absence of two adenines at position 81 (Supplementary Fig. 2b)

Guide: PheSF_guide_New_SV / PheSR_guide_New_SV

Repair template: lift_NDH2_aldo3utr_F_new / lift_NDH2_3xHA_R

### Cgd8_380 in situ HA-epitope

The insert encodes for a triple HA epitope followed by Nluc and the neomycin phosphotransferase drug-selection marker. We also included a reverse COWP1 3’UTR at the 3’ end of the repair to assure correct expression of the next downstream gene which is transcribed on the minus strand (Supplementary Fig. 2b).

Guide: 8_380_guide_F/8_380_guide_R

Repair template: 8_380_tag_F_NEW/8_380_tag_R_NEW

### Cell culture and *Cryptosporidium* infections

HCT-8 cells were purchased from ATCC (CCL-224TM) and maintained in RPMI 1640 medium (Sigma-Aldrich) supplemented with 10% Cosmic calf serum (HyClone) at 37°C in presence of 5% CO_2_. Oocysts were treated with 10 mM HCl at 37°C for 45-60 minutes before washing and resuspension in medium containing 1% serum, 0.2 mM sodium taurocholate, and 20mM sodium bicarbonate (infection media) to induce excystation. Infection media containing oocysts were transferred immediately onto cells and remained for the duration of the infection.

### Dose-response assay and IC50 calculations

In 96-well plates, HCT-8 cells were infected with 10,000 oocysts per well and incubated at 37°C for 3 hours. Equivalent volumes of clofazimine (2X final concentration) or 0.5% DMSO in infection media were added to the wells and incubated at 37°C for 48 hours. Medium was aspirated, cells were lysed and mixed with NanoGlo substrate (Promega), and luminescence was measured using a Glomax reader (Promega) (REF). IC50 values were calculated in GraphPad Prism software v9 (at least two independent experiments, each conducted with 5 replicates).

### Mouse infections and clofazimine treatment

All mouse infections were performed using 4- to 8-week-old male and female Ifnγ^−/−^ (Jackson Laboratory stock no. 002287) mice bred in-house. Mice were pretreated with antibiotic water and infected via oral gavage as detailed in^28,71^. Clofazimine (Sigma-Aldrich cat no. C8895) was formulated in MC-Tween (0.5% methylcellulose and 0.5% Tween-80) or PEG-Glucose (70% polyethylene glycol 400 and 1.5% glucose) suspension and given to mice orally. Feces of clofazimine- and vehicle-treated mice were collected and pooled as shown in Fig. 1F, 1G.

### Oocyst purification and genomic DNA extraction

Fecal material of infected mice was collected and oocyst were purified using a sucrose gradient and CsCl flotation ^71^. Genomic DNA was extracted using phenol/chloroform as described ^29^.

### Library preparation and Illumina sequencing of genomic DNA from cross progeny

We prepared Illumina libraries from extracted genomic DNA and sequenced both parents and 6 segregant pools. The library preparation was carried out using Illumina DNA Prep (former Nextera DNA Flex kit, Illumina Inc.). Subsequently, sequencing was performed on the Illumina NextSeq 2000 sequencer, utilizing P2 300 cycle flowcell kit.

### Genotype calling

The recent Iowa II telomere to telomere *de novo* assembly ^72^ was used as reference genome to identify single-nucleotide polymorphism (SNP) distinguishing the two parents, which were then used for bulk segregant analysis. Whole-genome sequencing reads for each library were individually mapped to the Iowa II *de novo* assembly using the BWA-MEM alignment algorithm with default parameters^73^. The resulting alignments were converted to SAM format, sorted into BAM format, and deduplicated using Picard tools^8^. Variants for each sample were called using HaplotypeCaller from GATK Suite and were subsequently aggregated across all samples using GenotypeGVCFs^74^ (see https://github.com/ruicatxiao/Automated_Bulk_Segregant_Analysis for detailed parameters).

### Bulk segregant analysis

SNP loci with coverage below 30× in either of the compared pools were excluded from bulk segregant analysis. At each variable locus, we counted reads corresponding to the genotypes of each parent and calculated allele frequencies. Iowa II allele frequencies were plotted across the genome, and outliers were removed using Hampel’s rule with a window size of 100 loci. Bulk segregant analysis was performed using the R package *QTLseqr* ^75^. Extreme QTLs were defined as loci with false discovery rates (FDRs, Benjamini-Hochberg adjusted *p*-values) below 0.01. Summary of the analysis and all metadata can be found in Supplementary Data 2.

### cgd7_1G00 ΔAA allele frequency analysis using publicly available sequence read archives

We have developed a Python pipeline, sra2vcf (https://github.com/ruicatxiao/sra2vcf), that performs robust and comprehensive SNP/INDEL analysis on both local and online SRA datasets for long-read or short-read DNA/RNA sequencing. Briefly for each sample the pipeline use sra-toolkit to download SRA and BWA is used to map reads to the genome. Mapped reads are sorted by genome coordinates using SAMTools, then duplicated reads are marked and removed with GATK suite. Bcftools mpileup is used to call variants using an INDEL detection optimized illumina-1.20 model. The output vcf are filtered by variant coverage and quality. Individual SRA sample’s vcf are aggregated to generate the final output table. The pipeline can accumulate and add new samples without reprocessing existing ones, and it intelligently checks for existing intermediate outputs to avoid redundant computations.

### High impact variant analysis for essential and non-essential genes using publicly available SRAs

The sra2vcf pipeline generates individual VCF files that are first processed with SnpEff to annotate variant effects on protein coding frames for each sample, followed by SnpSift aggregation to identify HIGH impact variants shared across samples. We developed Python programs goi_af.py and goi_cov.py to analyze SnpEff-annotated VCF files along with a gene-of-interest list. The output reports allele frequencies and allele coverage for each HIGH impact variant. We divide each gene’s coding regions into 10 bins from 5’ to 3’ to generate comprehensive allele frequency tables for target genes of varying length. All analysis codes are publicly available at https://github.com/ruicatxiao/cparvum_ndh2_clofazimine-resistance.

### Nanopore sequencing of amplified genomic material from cell cultures and fecal samples

Parasite genomic DNA was extracted from either cell culture supernatants or fecal samples as described in^29^. PCR amplification of the NDH2 locus was performed using PrimeStar Max ver. 2 (fwd primer: 5’-TCAAGTGGGGTCTCGGATG-3’; rev primer: 5’-CCCCACCCAGTACCTAAGATG-3’), then purified using Bioneer AccuPrep Gel/PCR Purification kit before sequencing. Nanopore sequencing of purified PCR amplicons was conducted using a commercial service provided by Eurofins Genomics.

### Genomic sequencing analyses

Raw read fastq files were mapped to the BG Iowa II *de novo* assembly using the BWA-MEM alignment algorithm with default parameters^73^. The resulting alignments were converted to SAM format, sorted into BAM format, and deduplicated using Picard tools^8^. Alignments were visualized in IGV to calculate deletion frequencies. All codes used for this analysis are available through github repository https://github.com/ruicatxiao/cparvum_ndh2_clofazimine-resistance.

### Engineering transgenic strains

Guide oligonucleotides (Sigma-Aldrich) were introduced into the *C. parvum* Cas9/U6 plasmid by restriction cloning and repair templates were constructed by Gibson assembly (New England Biolabs) ^71^ and excysted sporozoites were transfected as described ^71^. Briefly, 1.56 × 10^7^ *Cp* BG Iowa II oocysts or 5 x 10^6^ *Cp* KVI oocysts were incubated at 37 °C for 1 hr in 10 mM HCl followed by two washes with phosphate buffered saline (PBS) and an incubation at 37 °C for 1 h in 0.2 mM sodium taurocholate and 20 mM sodium bicarbonate to induce excystation ^29^. Excysted sporozoites were electroporated and used to infect mice as described ^29^. Integration was validated by PCR mapping and/or Sanger sequencing.

### Immunofluorescence assay

HCT-8 cells were seeded on coverslips in 24-well plates prior to infection. Infected coverslips were fixed with 4% paraformaldehyde (PFA; Sigma-Aldrich) in PBS for 20 min and then permeabilized with 0.25% Triton X in PBS for 10 min at room temperature. Coverslips were then blocked with 1% bovine serum albumin (BSA) in PBS for 1 hr before primary antibody (1:1000 rat anti-HA,1:500 rabbit anti-IMC3 or 1:1000 biotinylated anti-VVL) incubation followed by secondary antibody ( 1:1000 Alexa Fluor 488 anti-rat, 1:1000 Alexa Fluor 594 anti-rabbit or 1:1000 Alexa Fluor 594 streptavidin) incubation, both for 1 hr in 1% BSA. Coverslips were mounted using fluoro-gel mounting medium (Electron Microscopy Sciences) and imaged using the widefield Leica DM6000B or GE Delta Vision OMX.

### Ultrastructure expansion microscopy

Ultrastructure expansion microscopy was applied to *Cryptosporidium* infected HCT-8 cells as described for sporozoites ^58^. Infected coverslips were fixed in 4% paraformaldehyde for 20 min at 25°C and washed thrice with PBS before incubating overnight at 37°C in acrylamide and formaldehyde to prevent protein crosslinking. Samples were embedded in a water-based gel then denatured at 95°C prior to expansion in water to 4-5X their original size. Gels were shrunk in PBS before blocking and staining to save reagents. After re-expansion in water, the gels were imaged using the Leica Stellaris FALCON confocal microscope.

### Biochemical assays

Recombinant CpNDH2 was expressed as a maltose-binding protein (MBP) fusion using the pMAL-c5X vector (New England Biolabs, Ipswich, MA, USA). Clofazimine (purity ≥99%) and NADH (purity ≥98%) were purchased from Yuanye Bio-Technology (Shanghai, China). Menadione (purity ≥99.5%) was obtained from MedChemExpress (New Jersey, USA). Ubiquinone-2 (CoQ2; purity ≥95%) and menaquinone-4 (MK4; purity ≥98%) were purchased from GlpBio (Shanghai, China).

For cloning of the CpNDH2 gene (gene ID: cgd7_1900), the full-length open reading frame encoding wild-type CpNDH2 (CpNDH2-WT) was amplified from *C. parvum* genomic DNA (*GpC0* subtype IIdA19G1) by PCR using Phanta Max Super-Fidelity DNA Polymerase (Vazyme International, Nanjing, Jiangsu, China). The primers used were CpNDH2-forward (5′-cgcgatatcgtcgacggatccATGTCTAACTCTGAAAAGAATACTTCCAA-3′) and CpNDH2-reverse (5′-agcttatttaattacctgcagTTAGTGAGAAACGTTCATTTTGTAGATT-3′). Lowercase letters indicate additional sequences included for seamless cloning. The PCR amplicon was assembled into pMAL-c5X using the LightNing DNA Assembly Mix Plus kit (BestEnzymes Biotech, Lianyungang, Jiangsu, China). Recombinant plasmids were propagated in *E. coli* TOP10, purified, and sequence-verified for correct insertion.

For protein expression, the verified construct was transformed into *E. coli* BL21(DE3). Cultures were grown at 37 °C to an OD_450_ of ∼0.6, then induced with 0.5 mM IPTG at 25 °C for 6 h. Cells were harvested, lysed by sonication, and recombinant proteins purified using amylose-resin columns according to the manufacturer’s instructions (New England Biolabs). The quality and yield of purified proteins were assessed by SDS-PAGE and Bradford assay, using bovine serum albumin as the standard.

Tag-free CpNDH2 was prepared by cleavage of MBP-CpNDH2 with factor Xa protease (New England Biolabs). Reactions were carried out at room temperature for 24 h in 200 μL of buffer containing 20 mM Tris (pH 8.0), 100 mM NaCl, 2 mM CaCl₂, 200 μg MBP-CpNDH2, and 4 μg factor Xa. Following cleavage, proteins were dialyzed against buffer (50 mM Tris, pH 8.0, 150 mM NaCl, 1 mM EDTA) at 4 °C for 12 h, and the MBP tag was removed by amylose-resin purification. MBP alone was similarly expressed and purified for use as a negative control and for background subtraction in assays.

The catalytic activity of CpNDH2-WT was measured using a spectrophotometric assay adapted from published protocols^34–36^. Reactions were carried out at 25 °C in 100 μL of buffer containing 50 mM Tris (pH 7.0), 1 mM EDTA, 0.1% Triton X-100, 400 μM NADH, 4 μM CpNDH2 (MBP-fusion or tagless), and substrate at the indicated concentrations. Reactions were initiated by substrate addition, and NADH oxidation was monitored at OD_340_ in a microplate reader (BioTek Instruments, Santa Clara, CA, USA) at 0.5-min intervals for up to 40 min.

Since MBP-tagged and tag-free CpNDH2 displayed comparable activity toward menadione (MD) (Supplemental Fig. 1), subsequent assays with MK4, CoQ2, and clofazimine (CFZ) were performed using intact MBP-CpNDH2-WT. Optical density data were plotted against substrate concentrations, and kinetic parameters were estimated by nonlinear regression using a sigmoidal curve fit that incorporated the Hill coefficient. This yielded *K*_0.5_ (or *K*′) and *V*_max_ values. When the Hill slope was close to 1.0, the reaction followed Michaelis–Menten kinetics and *K*_0.5_ was equivalent to *K*_m_.

### Western blotting

Western blot was performed using rabbit anti-HA antibody diluted 1/1000 and mouse anti-H3pan antibody diluted 1/1000 and secondary anti-rabbit IRDye 800 and anti-mouse IRDye 680 diluted 1/10000. We followed the Licor best practice protocol. Blots were imaged using an Odyssey Licor device.

### Flow cytometry

1 million purified oocysts were washed and resuspended in 200 μL FACS buffer (1× PBS, 0.2% bovine serum albumin, 1 mM EDTA). Oocysts were then passed through a 70 μM mesh filter. Data were collected on a FACSymphony A3 Lite (BD Biosciences) and analyzed with FlowJo v10 software (TreeStar). Construct integration frequency was measured by positivity for an mNeonGreen reporter.

## Acknowledgements

This work was supported in part by grants from the National Institutes of Health to B.S. (R01AI112427, R01AI127798), B.S. and Christopher Hunter (R01AI148249), to the Penn Vet Imaging Core (S10OD021633), Swiss National Science Foundation fellowships 402 P2BEP3_191774 and P500PB_211097 to S.S, and European Molecular Biology Organization fellowship ALTF1145-2021 to A.B. We thank Adam Sateriale for initial contributions, Jessica Byerly and Chloe Tang for support of animal experimentation, and Bethan Wallbank, Allison Cohen, Abigail Daniels, Chloe Tang, and Maria Merolle for sharing transgenic parasites.

## Contributions

Conceptualization: B.S., G.Y.B., S.S.

Methodology: A.B., K.M.O., P.J., B.X., D.W., R.X., G.Y.B., S.S.

Investigation: A.B. (expansion microscopy, Fig. 5a–b); K.M.O. (flow cytometry, Fig. 4g); P.J., B.X., D.W. (biochemical assays, Fig. 4i–k and Supplementary Fig. 1); R.X. (allele frequency calculations, Fig. 6a–c); G.Y.B., S.S. (all other experiments).

Formal analysis: R.X., G.Y.B., S.S., A.B., K.M.O., P.J., B.X., D.W.

Visualization: G.Y.B., S.S., B.S.

Writing original draft: G.Y.B., S.S., B.S.

Writing review C editing: All authors. Supervision: B.S.

## Data availability

Whole genome raw sequencing data and the raw amplicon sequencing data have been deposited in the NCBI’s Sequencing Read Archive database under Bioproject numbers PRJNA1336748 and PRJNA1337473, respectively.

## Supplementary Information

**Supplementary Figure 1:**
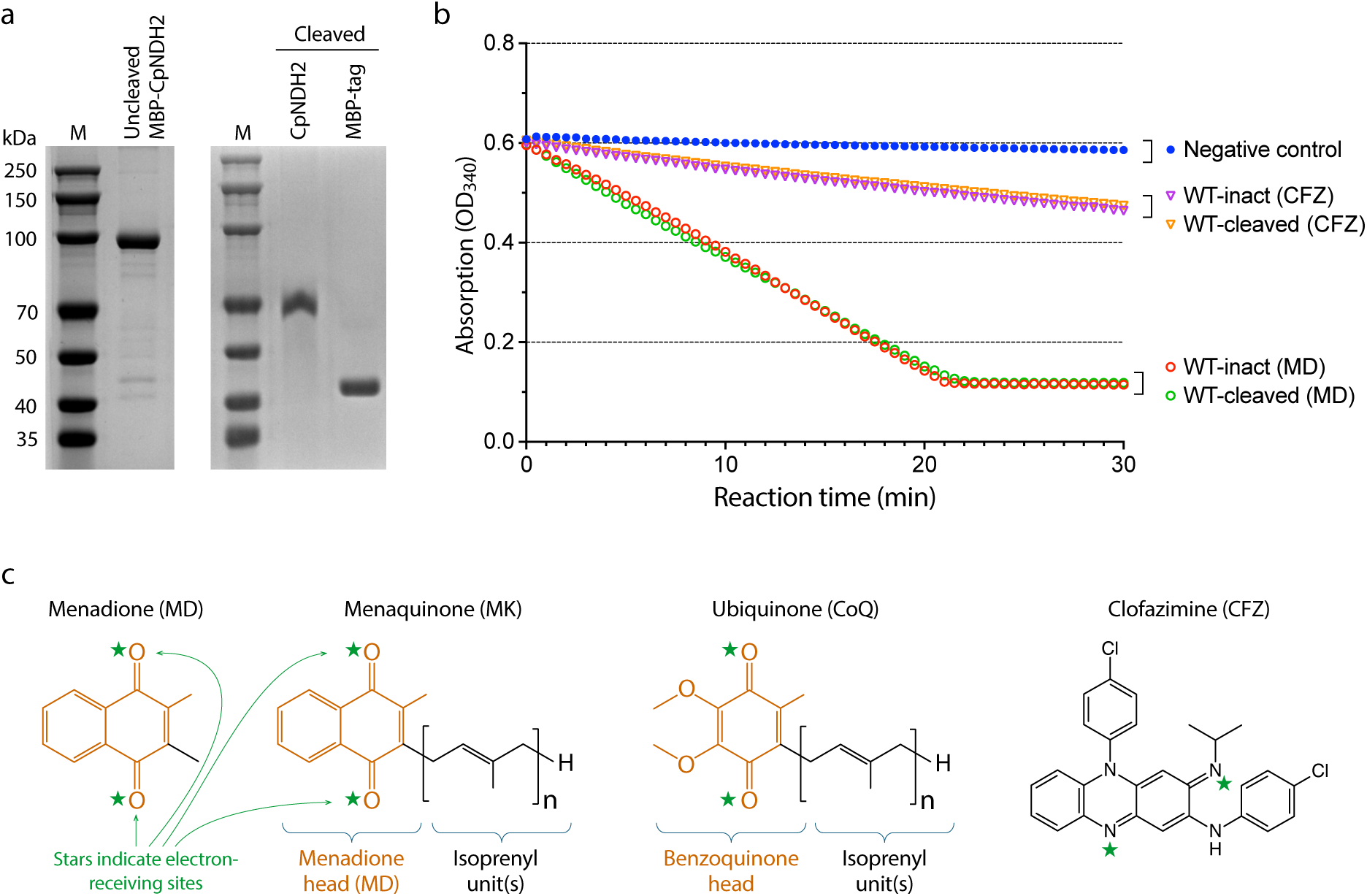
Initial assessment of catalytic activity using purified recombinant wild-type CpNDH2 protein in intact and cleaved forms. **(a)** SDS-PAGE gels showing purified intact (uncleaved) and cleaved MBP-CpNDH2 protein. **(b)** Initial assessment of intact and cleaved CpNDH2 proteins in catalyzing the electron transfer from NADH to menadione (MD) and clofazimine (CFZ) using a spectrometeric assay.**(c)** Chemical structures of menadione, menaquinone, ubiquinone, and clofazimine. Green stars indicate atoms positioned to accept electrons during enzymatic reduction.

**Supplementary Figure 2:**
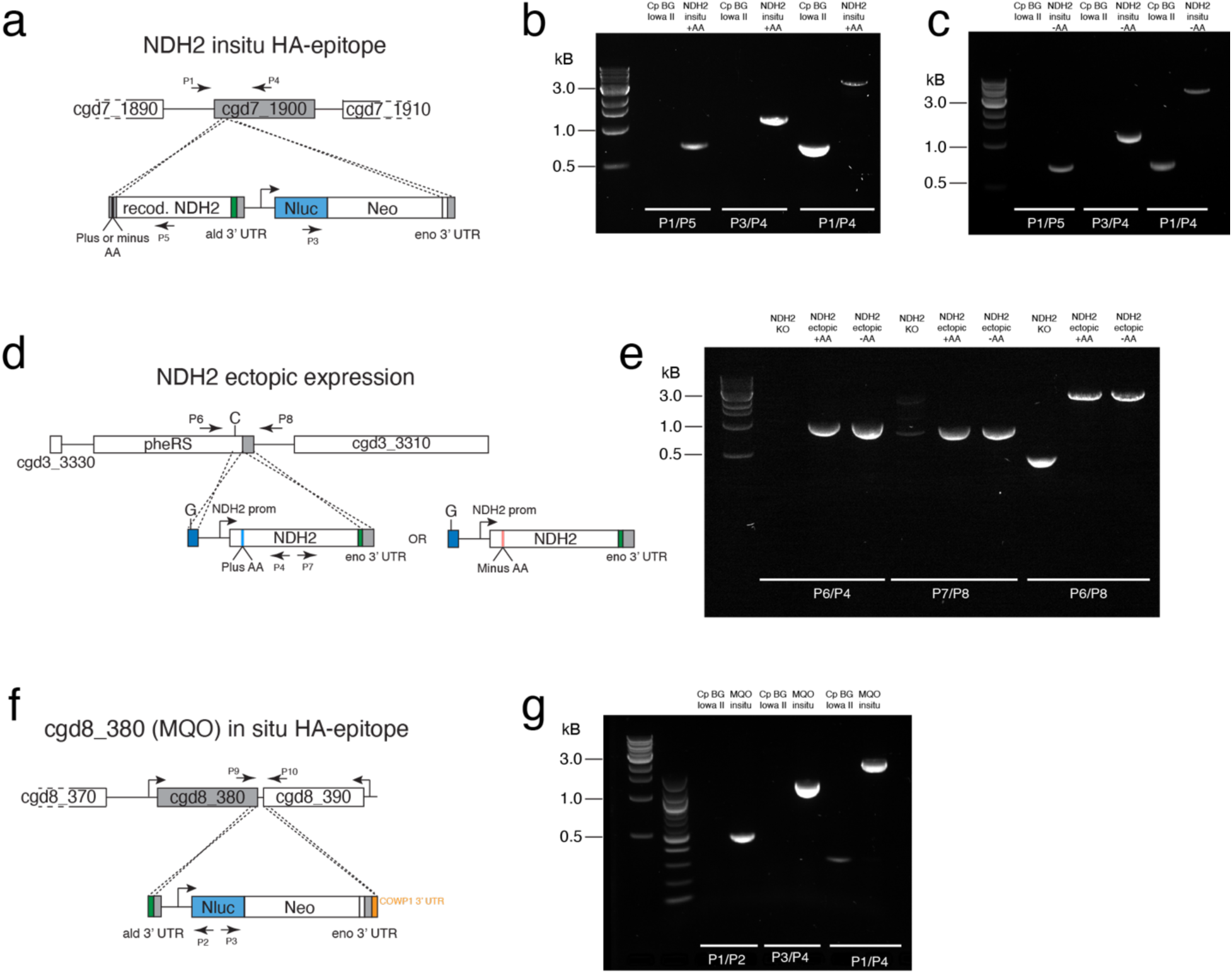
Transgenic parasite strains constructed for this study. (**a**) Map of NDH2 insitu HA epitope tagging strategy. Two strains were generated, either with or without INDEL (plus or minus AA). HA epitope = green. (**b**) Gel shows PCR mapping of the plus AA strain described in (a). (**c**) Gel shows PCR mapping of the minus AA strain described in (a). (**d**) Ectopic expression of NDH2-HA and NDH2-ΔAA-HA. HA epitope = green. (**e**) Gel shows PCR mapping of the strains described in (d). (d). (**f**) Map of MQO insitu HA epitope tagging strategy. HA epitope = green. (**g**) Gel shows PCR mapping of the MQO insitu HA epitope strain. Labelling of the gel refers to the amplicons shown in the maps (f). Labelling of all the gels refer to the amplicons shown in the respective maps.

